# Cold exposure induces dynamic changes in circulating triacylglycerol species, which is dependent on intracellular lipolysis: a randomized cross-over trial

**DOI:** 10.1101/2022.09.09.507312

**Authors:** Maaike E. Straat, Lucas Jurado-Fasoli, Zhixiong Ying, Kimberly J. Nahon, Laura G.M. Janssen, Mariëtte R. Boon, Gernot F. Grabner, Sander Kooijman, Robert Zimmermann, Martin Giera, Patrick C.N. Rensen, Borja Martinez-Tellez

## Abstract

**Background:** The application of cold exposure has emerged as an approach to enhance whole-body lipid catabolism. The global effect of cold exposure on the lipidome in humans has been reported with mixed results depending on intensity and duration of cold.

**Methods:** This secondary study was based on data from a previous randomized cross-over trial (ClinicalTrials.gov ID: NCT03012113). We performed sequential lipidomic profiling in serum during 120 min cold exposure of human volunteers. Next, the intracellular lipolysis was blocked in mice using a small-molecule inhibitor of adipose triglyceride lipase (ATGL; Atglistatin), and were exposed to cold for a similar duration. The quantitative lipidomic profiling was assessed in-depth using the Lipidyzer platform.

**Findings:** Cold exposure gradually increased circulating free fatty acids reaching a maximum at 60 min, and transiently decreased total triacylglycerols (TAGs) only at 30 min. A broad range of TAG species was initially decreased, in particular unsaturated and polyunsaturated TAG species with ≤5 double bonds, while after 120 min a significant increase was observed for polyunsaturated TAG species with ≥6 double bonds. The mechanistic study in mice revealed that the cold-induced increase in polyunsaturated TAGs was largely prevented by blocking adipose triglyceride lipase.

**Interpretation:** We interpret these findings as that cold exposure feeds thermogenic tissues with TAG-derived fatty acids for combustion, resulting in a decrease of circulating TAG species, followed by increased hepatic production of polyunsaturated TAG species induced by liberation of free fatty acids stemming from adipose tissue.

**Research in Context:** *Evidence before this study:* Cold exposure has emerged as a novel non-pharmacological strategy to enhance whole-body lipid catabolism, to improve lipid homeostasis and ultimately cardiometabolic health. In mice, cold exposure accelerates the clearance of triacylglycerol (TAG)-rich lipoproteins from the circulation, reducing circulating TAGs. In humans however, the effect of cold exposure on whole-body TAG metabolism remained thus far controversial, as mixed results are reported depending on intensity and duration of cold.

*Added value of this study:* Here, we performed sequential lipidomic profiling in serum during 120 min cold exposure of human volunteers. We show that cold exposure gradually increases circulating free fatty acids and transiently decreases total TAGs after 30 min, accompanied with a dynamic change in circulating TAGs as dependent on their saturation status and length. Specifically, cold exposure decreases a broad range of more saturated and shorter TAG species, while after 90-120 min polyunsaturated TAG species with ≥6 double bonds start to increase. Subsequently, we performed a mechanistic study in mice, in which we show that the cold-induced increase in polyunsaturated TAGs is largely prevented when blocking intracellular lipolysis.

*Implications of all the available evidence:* Our findings describe a mechanism by which cold exposure provides thermogenic tissues with TAG-derived fatty acids for combustion. At the same time, cold exposure increases lipolysis in white fat to drive hepatic TAG production to further feed thermogenic tissues. For the first time, these results show that the TAG lowering effect of cold exposure as observed in mice can be recapitulated in humans, which warrants further studies on the beneficial effects of cold exposure on accelerating lipid metabolism to improve cardiometabolic health.

## Introduction

Worldwide, the prevalence of adiposity and its associated cardiometabolic diseases (*e*.*g*., type 2 diabetes mellitus, fatty liver disease, and cardiovascular diseases) is increasing at an alarming rate and has become a major public health concern^1,2^. One of the contributing factors in the development of cardiometabolic diseases is dysfunctional lipid homeostasis arising from impaired storage, mobilization, and/or combustion of lipids by metabolic organs (*e*.*g*., white adipose tissue, liver, skeletal muscle, brown adipose tissue)^3^. Over the last decade, increased interest has emerged for non-pharmacological strategies to restore lipid homeostasis, among which the application of cold exposure to stimulate non-shivering thermogenesis.

Physiologically, cold exposure stimulates the sympathetic nervous system to release norepinephrine that binds to β-adrenergic receptors on thermogenic tissues^4^. In mice, brown adipose tissue is the most important thermogenic tissue during non-shivering thermogenesis. Cold exposure enhances the translocation, and therewith activity, of lipoprotein lipase to the cellular membrane of brown adipocytes, which involves reduced expression of angiopoietin-like 4 (Angptl4)^5^. This allows increased uptake of triacylglycerol (TAG)-rich lipoprotein-derived fatty acids (FAs) via CD36/FATP that are catabolized for heat production^6,7^. Therewith, cold exposure in mice accelerates the clearance of TAG-rich lipoproteins from the circulation, resulting in reduced circulating TAGs after cold exposure^6-10^. Cold-activated brown adipose tissue in mice thus has the potential to alleviate hyperlipidemia, which is beneficial in pathological conditions such as obesity or genetic hypertriglyceridemia^6,8^. In addition, activation of brown adipose tissue in mice indirectly attenuates hypercholesterolemia, due to increased hepatic uptake of the delipidated, cholesterol-enriched lipoprotein remnants, therewith protecting from atherosclerosis development^8,10^.

The question remains to what extent the impact of cold exposure on TAG-rich lipoprotein metabolism translates to humans. It has been shown with positron emission tomography-computed tomography (PET-CT) analysis that short-term cold stimulation increases the uptake of free FAs (FFA) by brown adipose tissue^11-13^, but a PET-compatible tracer to study cold-induced uptake of TAG-derived FAs by thermogenic tissues is not available as of yet. Hence, the effect of cold exposure on whole-body TAG metabolism is less clear, especially as the total circulating TAG level in response to cold does not change^11,14^, or even increases^15,16^ dependent on the intensity and duration of the cold exposure. In turn, the aim of this study was to obtain more insight in the effects of cold exposure on whole body TAG metabolism in humans by assessing sequential effects of short-term cold exposure (*i*.*e*., 30, 60, 90, and 120 minutes) on the serum lipidome of healthy men through comprehensive, quantitative in-depth lipidomic profiling using the Lipidyzer platform^17^. To obtain mechanistic insight into the role of intracellular lipolysis and its relation to lipidomic changes during cold exposure, we blocked intracellular lipolysis in mice using a small-molecule inhibitor of adipose triglyceride lipase (ATGL; Atglistatin^18^) and exposed them to cold for a similar duration.

## Methods

### Human experiment

#### Study participants

A total of 10 young (age: 24.4 ± 1.0 years), healthy, lean (body mass index: 22.7 ± 0.6 kg/m^2^) Dutch Europid men were included in this study^19^. The inclusion criteria were as follows: (i) male; (ii) age between 18 and 30 years; (iii) body mass index <25 kg/m^2^. Exclusion criteria included: (i) presence of any chronic endocrine or cardiovascular disease as determined by medical history and physical examination; (ii) the use of drugs that is known to influence the cardiovascular or endocrine system, or the use of drugs that is known to influence glucose or lipid metabolism; (iii) smoking or abuse of alcohol or drugs; (iv) excessive weight loss or exercise. Detailed study subject characteristics have been previously described^19^.

#### Study design

This study was a secondary analysis of a randomized crossover trial, with three different treatment regimes. In this study the metabolic effects of mirabegron ingestion were compared with cold exposure, using a personalized cooling protocol, and with placebo in a control experiment, in a randomized double-blind controlled fashion. In the current secondary analysis only data from cold exposure and the control experiment at room temperature are included (**Figure 1A**). The randomization and allocation have been extensively described elsewhere ^19^. The cooling protocol took place from 10:30 AM till 12:30 PM at the Leiden University Medical Center (LUMC, The Netherlands), the control experiment took place from 9:00 AM till 11:00 AM. The evening prior to the experiments, a standardized meal was consumed after which participants remained fasted from 10:00 PM overnight until the end of the experiments (10h). In addition, participants were asked to restrict from exercise 48 hours in advance and to not consume alcohol or caffeine containing beverages 24 hours in advance. During all the experiments, participants stayed in a clinical examination room and lied down in a bed in semi-supine position.

**Figure 1.**
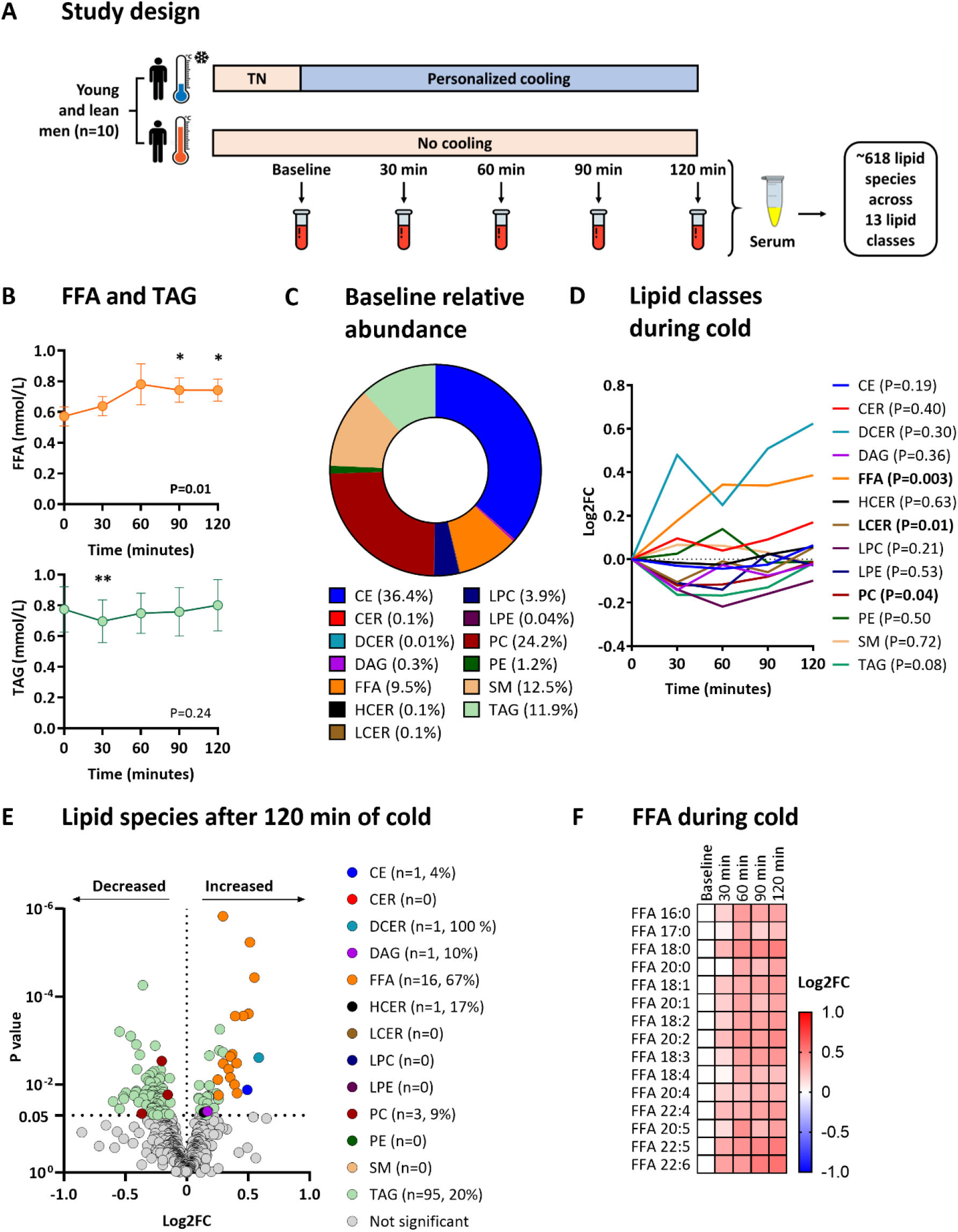
Study design and lipidomic changes during cold exposure. (**A**) Overview of the crossover study design. During the first occasion, participants were exposed to a personalized cooling protocol. During the other occasion, participants were exposed to constant room temperature. Under both conditions, venous blood was drawn at five sequential timepoints (*i*.*e*., baseline, 30, 60, 90, and 120 min) to obtain serum for longitudinal lipidomic profiling. (**B**) Dynamic changes in total free fatty acid and triacylglycerol concentration during cold exposure, as measured with enzymatic assays. Data is presented as mean and standard error of the mean. P values are obtained from ANOVA repeated measures. *P<0.05, **P<0.01; compared to baseline (*i*.*e*., 0 min). (**C**) Baseline relative abundance of the number of lipid classes. (**D**) Dynamic changes in lipid classes during cold exposure. Data is presented as the log2 fold change (log2FC) relative to baseline (*i*.*e*., 0 min). P values are obtained from ANOVA repeated measures. (**E**) Volcano plot showing the change of individual lipid species after 120 min of cold exposure. The X-axis shows the log2FC between 120 min of cold exposure *vs*. baseline, the Y-axis shows the P value. P values are obtained from paired Student’s t test. Values between brackets indicate the absolute number and the percentage of lipid species within the lipid class that were modulated by cold exposure. (**F**) Heatmap of free fatty acid (FFA) species that significantly dynamically changed during cold exposure. The color of the squares represents the log2FC relative to baseline. FFAs are sorted by the number of double bonds. ANOVA repeated measures were used to compare the different timepoints. CE, cholesteryl esters; CER, ceramides; DAG, diacylglycerols; DCER, dihydroceramides; FFA, free fatty acids; FC, fold change; HCER, hexosylceramides; LCER, lactosylceramides; LPC, lysophosphatidylcholine; LPE, lysophosphatidylethanolamine; PC, phosphatidylcholine; PE, phosphatidylethanolamine; SM, sphingomyelin; TAG, triacylglycerols; TN, thermoneutral.

At the start of the first experimental session, eligibility of participants was assessed during a screening procedure, that included a medical history questionnaire, anthropometric measurements, analysis of their body composition (Bodystat 1500, Bodystat, UK). An intravenous cannula was inserted in the antecubital vein. Then, participants in underwear were exposed to a personalized cooling protocol using two water-perfused cooling mattresses (BlanketRol^®^ III, Cincinnati Sub-Zero Products, USA) (**Figure 1A**). Initially, water temperature was set at 32°C, which is considered thermoneutral, as baseline period for 1 hour. After that, the water temperature gradually decreased by 5°C every 10 minutes, until the participants started to shiver. Shivering was specified as unstoppable contraction of the muscles and was both reported by the participants and confirmed visually by researchers. At this point, temperature was raised by 3°C to stop shivering and ensure maximal non-shivering thermogenesis. Then, the stable cooling period of 60 minutes started. During the personalized and stable cooling period, blood samples were obtained at 0 minutes, 30 minutes, 60 minutes, 90 minutes, and 120 minutes. During the control experiment, blood sampling was carried out at room temperature (21°C) in the same participants but wearing hospital clothing instead. No important changes were performed in the methodology or outcomes after the beginning of trial.

#### Blood collection

Blood was collected in SST gel tubes which were centrifuged for 10 min at 2200*g* to obtain serum. These serum samples were aliquoted and stored at -80°C until further batch-wise analyses as previously described^19^. Commercially available enzymatic kits were used to measure serum concentrations of TAGs (Roche Diagnostics, Woerden, the Netherlands) and FFAs (Wako chemicals, Nuess, Germany).

### Animal experiment

A total amount of eighteen 10-week-old male wild-type mice (C57BL/6J background, Charles River Laboratories) were group-housed at 30°C for 4 days, whereafter they were randomized to one of the 3 treatment groups, namely: control mice, cold exposed mice, and cold exposed mice treated with Atglistatin (RandoMice software, Leiden, The Netherlands^20^). After that, they were individually housed in metabolic home-cages (Promethion Line, Sable Systems, Las Vegas, USA) under thermoneutral temperature (30°C) and 12:12 hour light-dark cycle with *ad libitum* access to water and standard laboratory chow diet (Rat and Mouse No. 3 Breeding, SDS, Horley, United Kingdom) for 3 days. At day 8, all animals were fasted for 4 hours (8 AM - 12 PM) after which they received an intraperitoneal injection with vehicle (100 µL of phosphate buffered saline [PBS] containing 0.25 % Cremophor EL; control mice and cold exposed mice) or the small-molecule ATGL inhibitor Atglistatin (5 μmol in 100 µL vehicle; cold exposed mice treated with Atglistatin). Just after injection, control mice (n=6) remained at 30°C and cold exposed mice with (n=6) or without (n=6) Atglistatin treatment were gradually exposed to 10°C for the next 150 minutes. We extended the cold exposure to 150 min to mimic the human cooling conditions, as cooling the air temperature of metabolic cages takes longer. All animals remained fasted for this period with access to water. Afterwards, blood was drawn via a tail vein incision (70 µL) from every single animal. Plasma was collected and measured for total TAGs concentration using commercial enzymatic kits from Roche Diagnostics (10166588130; Mannheim, Germany). 25 µL of plasma were stored for the subsequent lipidome analyses.

### Quantitative lipidomics using the Lipidyzer platform

#### Sample preparation

Serum (humans) and plasma (mice) lipids were extracted using methyl-tert-butylether in a randomized order as previously described^17^. To 25 µL of the sample, the following was added: 160 µL of methanol, 50 µL of internal standard solution (Lipidyzer internal standard kit, [cat# 5040156, Sciex, Redwood City, CA, USA] containing >50 labelled internal standards for 13 lipid classes), and 550 µL of methyl-tert-butylether. Samples were vortexed at room temperature for 30 min. Thereafter, 200 µL of water was added for phase separation and the samples were centrifuged at 13,100*g* for 10 min at 20°C. The upper layer was transferred to a glass vial and the lipid extraction was repeated adding 300 µL of methyl-tert-butylether, 100 µL of methanol, and 100 µL of water. Subsequently, the organic extracts were combined and dried under a gentle stream of nitrogen. The Lipidyzer running buffer which consists of 250 µL 10 mM of ammonium acetate in 50:50 (v/v) methanol:dichloromethane was added, and the samples were transferred to a glass vial with insert for injection.

#### Data acquisition

Lipid extracts were analyzed with the Lipidyzer platform (Sciex, Redwood City, CA, USA)^17^, which is a flow-injection-based ion-mobility triple quadrupole system that comprises a Sciex 5500 QTrap equipped with a SelexION differential mobility spectrometry (DMS) interface coupled to a UHPLC system (Shimadzu Nexera series, Kyoto, Japan) used for injection and delivering buffer at 7 µL/min. All the lipids were measured in a randomized batch-controlled fashion. The samples were measured in 11 consecutive batches. Lipid species were identified and quantified using multiple reaction monitoring (MRM) and positive/negative ionization switching. A total of 50 µL of the resuspended samples were injected using 2 acquisition methods. Method #1 operated with active DMS separation under the following conditions: DMS temperature low, modifier (propanol) composition low, separation voltage 3500V, DMS resolution enhancement low. In Method #2, the DMS cell was not activated. The MS operated under the following conditions: curtain gas 17, CAD gas medium, ion spray voltage 4100V in ESI + mode and -2500 V in ESI-mode, temperature 200°C, nebulizing gas 17, and heater gas 25.

The following 13 lipid classes were quantitatively assessed: cholesterol ester (CE), ceramides (CER), diacylglycerides (DAG), dihydroceramides (DCER), FFAs, hexosylceramides (HCER), lysophosphatidylcholine (LPC), lysophosphatidylethanolamine (LPE), phosphatidylcholine (PC), phosphatidylethanolamine (PE), sphingomyelin (SM), TAGs. Firstly, LPC, LPE, PC, PE, and SM lipid classes were analyzed applying method #1. Next, CE, CER, DAG, DCER, FFA, HCER, LCER and TAG lipids were analyzed applying method #2. A detailed technical description including the list of all monitored transitions and the detailed experimental setting can be found elsewhere^17^. Four quality controls (QC) consisting of a commercial freeze-dried serum/plasma reference were added to each measurement batch.

#### Data processing

Lipid data were reported by the Lipidomics Workflow Manager software (Sciex, Redwood City, CA), which calculates concentrations of each detected lipid as average intensity of the analyte MRM/average intensity of the most structurally similar deuterated internal standards [Lipidyzer internal standard kit, (cat# 5040156, Sciex, Redwood City, CA, USA)] MRM multiplied by its concentration. The Lipidyzer platform provides several readouts which are the following: lipid class concentration (nmol/mL), lipid species concentration (nmol/mL), and FA concentration (nmol/mL), the latter being the concentration of all lipids of a specific lipid class containing specific FA species. Relative abundance (%) for the aforementioned items is provided. Lipid species were excluded for the analyses if the relative standard deviation (RSD) of the QCs was >20% within each batch or >25% between batches. Lipids that were detected in less than two-third of the sample or were not observed in every single batch were discarded and considered as missing values.

#### Lipid nomenclature

The Lipidyzer platform cannot specify the exact stereospecific numbering-position of the FA side chains. Moreover, the output format of the Lipidyzer platform does not strictly adhere to the lipid short-hand notation previously described^21^. To facilitate data handling and comparison in this study we have kept this format, how the Lipidyzer format translates into the lipid short-hand annotation can be found elsewhere^17^. Additionally, it must be noted that for the TAG lipids, the Lipidyzer platform can define only one of the three FA side chains. Hence, a TAG lipid specified as, for example, TAG 54:6-FA 18:1 would refer to a TAG lipid with 54 carbons, 6 double bonds, and 1 side chain being FA 18:1. This also results in the fact that one concatenated TAG lipid can be reported with various combinations of fatty acyl tails (*e*.*g*., TAG 54:6-FA 16:0, TAG 54:6-FA 20:4, and so on). Consequently, this leads to an overestimation of the number of measured TAG species and TAG 54:6 will be reported three times, once for each acyl side chain. While this annotation is advantageous for fatty acid composition analysis, it overestimates the number of TAGs reported. When reporting the TAG lipid class concentration, the Lipidomics Workflow Manager software corrects for this by dividing the summed concentration of all TAG combinations by a factor of three^17^.

### Ethics statement

#### Human study

The experimental study was registered on ClinicalTrials.gov (NCT03012113). Written informed consent was obtained from all volunteers prior to participation. The study was carried out according to the principles of the revised Declaration of Helsinki (41) and was approved by the Medical Ethical Committee of the Leiden University Medical Center (LUMC), Leiden, the Netherlands.

#### Animal study

The animal experiment was carried out according to the Institute for Laboratory Animal Research Guide for the Care and Use of Laboratory Animals, and were approved by the National Committee for Animal Experiments (Protocol No. 11600202010187) and by the Ethics Committee on Animal Care and Experimentation of the Leiden University Medical Center (Protocol No. PE.21.002.015). All animal procedures were conformed the guidelines from Directive 2010/63/EU of the European Parliament on the protection of animal used for scientific purposes.

### Statistics

Statistical analyses were performed using IBM SPSS Statistics (version 26.0, IBM Corporation, Chicago, IL, USA) and GraphPad Prism was used to build figures (version 7.0, GraphPad Software, San Diego, CA, USA). This study was a secondary study of a previous randomized cross-over trial. Therefore, no specific power calculation was performed for the current study. Normality was tested using the Shapiro-Wilk test. None of the outcomes followed a normal distribution, after which all values were log2-transformed. To investigate the final effect of cold exposure on the lipidomic profile, paired Student’s t test between the concentrations at baseline *vs*. 120 minutes was performed. To investigate the dynamic effect of cold exposure on the total lipid concentration (*i*.*e*., FFA and TAG) and the lipidomic profile in humans, one-way repeated measures analyses of variance (ANOVA) with the variable ‘time’ (baseline, 30 minutes, 60 minutes, 90 minutes and 120 minutes) as within-subject factors was performed. The same statistical method was used for the control experiment. To investigate the effect of cold exposure on the lipidomic profile in mice, unpaired Student’s t test between cold exposed mice (either treated with vehicle or with Atglistatin) *vs*. mice kept at thermoneutrality was performed. A p-value < 0.05 was considered statistically significant. To investigate the effect of cold exposure on the total TAG concentration in mice, an one-way ANOVA was used. Most figures show log2 fold changes (log2FC) relative to baseline per single timepoint. These fold changes were calculated with the log2-transformed outcomes (*i*.*e*., log2FC 120 min = lipid concentration at 120 minutes (in log2) minus lipid concentration at baseline (in log2)). Values in text represent mean and standard error of the mean.

## Results

### Cold exposure modulates the concentration of circulating lipid species

We first determined total circulating FFA and TAG concentration at all timepoints using conventional enzymatic assays. Cold exposure gradually increased the total concentration of FFAs, reaching statistical significance at both 90 min (+30%; 0.57±0.06 *vs*. 0.74±0.08 mmol/L, P=0.02) and 120 min (+30%; 0.57±0.06 *vs*. 0.74±0.07 mmol/L, P=0.01; **Figure 1B**). In contrast, the total concentration of TAGs transiently decreased after 30 min of cold exposure (−9%; 0.77±0.15 *vs*. 0.70±0.14 mmol/L, P=0.008), after which total TAG concentrations returned to baseline despite ongoing cooling (**Figure 1B**).

To obtain more detailed insight in the effect of cold on whole body lipid metabolism, we next quantified 765 lipid species across 13 lipid classes using the Lipidyzer platform. Lipids were considered as missing values and excluded from further analyses if they were not detected at all five timepoints in ≥ two-third of the participants. After exclusion of 147 missing values, the dataset contained 618 lipid species. At baseline, the relative abundance of the number of FFAs was 9.5% and that of TAGs was 11.9% (**Figure 1C**). We first determined the effect of cold exposure on all 13 lipid classes. Cold exposure modulated the concentrations of FFAs (P=0.003) in addition to those of lactosylceramides (P=0.01) and phosphatidylcholines (P=0.04). Cold exposure did not affect the concentrations of TAGs (P=0.08), although the direction of the TAGs was the same as determined with the enzymatic assay with a dip after 30 min of cold exposure (**Figure 1D**).

Next, we assessed the effect of cold exposure on the 618 lipid species across the 13 lipid classes. When comparing the lipid species’ concentrations between 120 min of cold *vs*. baseline, 72/618 lipid species were either significantly increased or decreased (12%, P<0.05; **Figure 1E**). Cold exposure increased the concentrations of 1/23 cholesteryl esters (4%), 1/1 dihydroceramides (100%), 1/10 diacylglycerols (10%), and 1/6 hexosylceramides (17%); and decreased the concentrations of 3/33 phosphatidylcholines (9%, all P<0.05). Moreover, cold exposure increased the concentrations of 16/24 individual FFA lipid species (67%) and the concentrations of 22/468 individual TAG lipid species (5%), whereas the concentrations of 73/468 individual TAG lipid species were decreased (16%, all P<0.05; **Figure 1E**).

### Cold dynamically modulates TAG lipid species dependent on their saturation status and length

Although only 72/618 lipid species had different concentrations after cold exposure (**Figure 1E**), as many as 232/618 lipid species were dynamically modified during cold exposure (38%, P<0.05; not shown). The majority of these lipid species included FFAs (n=15; **Figure 1F**) and TAGs (n=206; **Figure 2A**). Therefore, we next specifically focused on the characteristics of those FFA and TAG species and how they dynamically responded during cold exposure.

**Figure 2.**
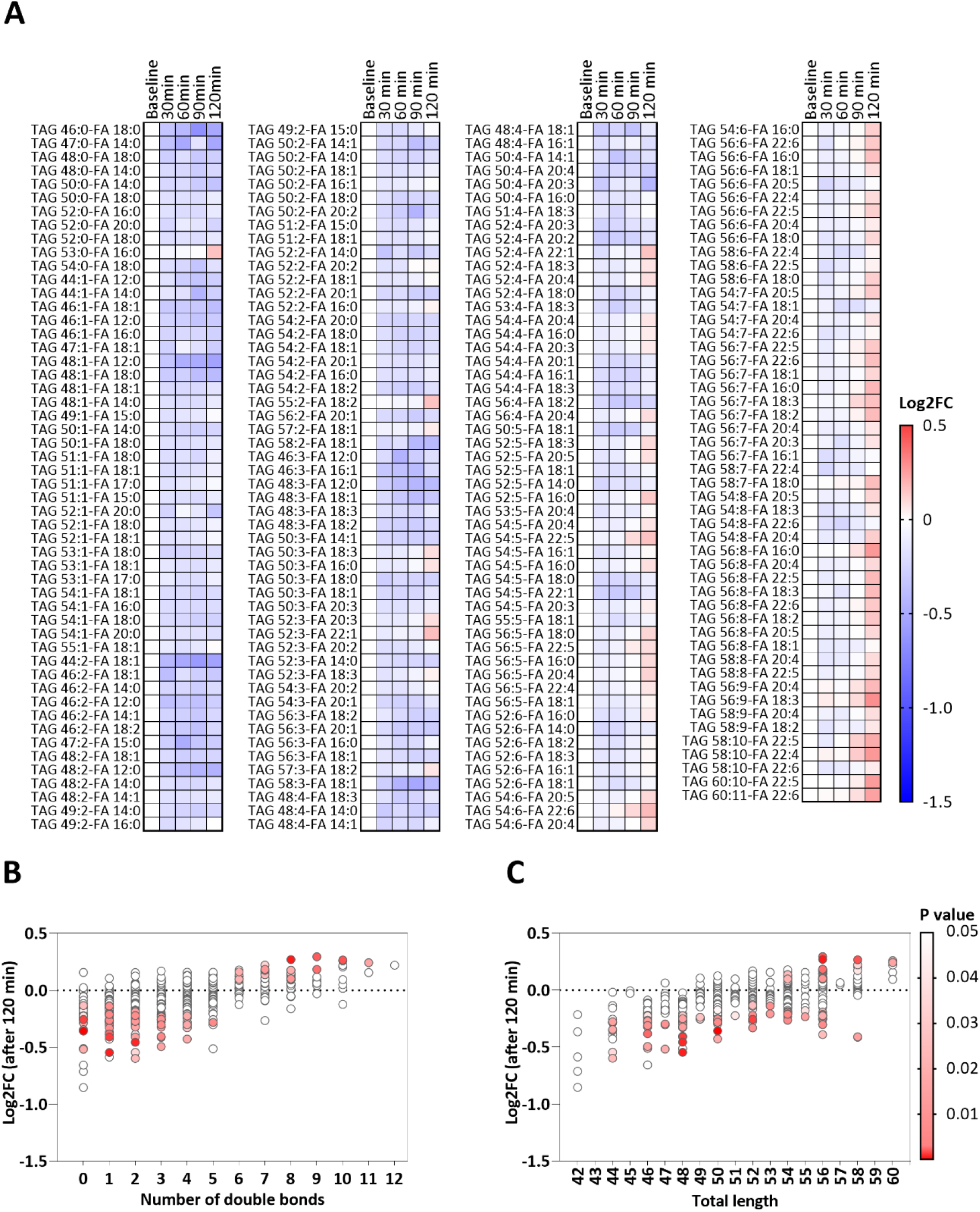
Triacylglycerol (TAG) species response to cold exposure. **(A)** Heatmap of TAG species that significantly dynamically changed during cold exposure. The color of the squares represents the log2 fold change (log2FC) relative to baseline. TAGs are sorted by the number of double bonds. ANOVA repeated measures were used to compare the different timepoints. **(B)** Bubble plot showing the P value (in color) and log2FC (on y-axis) after 120 minutes of cold exposure per TAG species categorized by number of double bonds. P values are obtained from paired Student’s t test. (**C**) Bubble plot showing the P value (in color) and log2FC (on y-axis) after 120 minutes of cold exposure per TAG species categorized by the total length of the TAG. P values are obtained from paired Student’s t test. FA, fatty acid; FC, fold change; TAG, triacylglycerol.

Concentrations of the FFA species (15/24, 63%) increased from the start generally until the end of cold exposure (**Figure 1F**). In contrast, the concentrations of the TAG species initially decreased upon cold exposure, followed by an increase of 88/468 TAG species (19%) after 90-120 minutes of cooling which was dependent on the saturation status of the TAGs (**Figure 2A**). Eventually, TAG species containing the lowest number of double bonds (*i*.*e*., ≤5 double bonds) were decreased, whereas TAG lipid species with a higher number of double bonds (*i*.*e*., ≥6 double bonds) were increased (**Figure 2B**). In addition, TAG species that were longer-chained (*i*.*e*., ≥56 carbons) were more likely to increase at the end of cooling (**Figure 2C**). Collectively, we observed that specifically levels of PUFA-containing TAGs increased at the end of cold exposure.

To elucidate whether the changes were solely an effect of cold exposure, we included a control condition in which participants were exposed to room temperature (21°C). Similar as during cold exposure, the total concentration of FFAs increased during the control experiment, leading to an increased concentration after 120 min (+38%; 0.32±0.03 *vs*. 0.44±0.06 mmol/L, P=0.049). The total concentration of TAGs did not change during the control experiment (P=0.47; **Supplementary figure 1A, B**). In addition, the concentrations of 9/24 FFA lipid species (38%) and 13/440 TAG lipid species (3%) were increased and 41/440 TAG lipid species (9%) were decreased (all P<0.05; **Supplementary figure 1C**). Importantly, during exposure to room temperature *vs*. cold, less FFA species (38 *vs*. 63%) and less TAG species (21 *vs*. 44%) were dynamically modulated (**Supplementary figures 1D, E** and **2**). When exposed to room temperature, TAG species with the lowest number of double bonds (*i*.*e*., ≤3 double bonds) generally decreased over time, whereas TAG species with a higher number of double bonds (*i*.*e*., ≥4 double bonds) generally increased over time (**Supplementary figure 2**).

### Intracellular lipolysis is critical for the cold-induced changes in the lipidome in mice

We hypothesized that the release of FFAs from the adipose tissue would drive the production of PUFA-containing TAGs by the liver, and that this process is accelerated during cold exposure. To test this hypothesis, we treated single-housed male C57BL/6J mice either with vehicle or with the ATGL inhibitor Atglistatin^18^, and exposed them for 150 min to a cold ambient temperature (10°C). The circulating lipidome was compared with control mice that were exposed to thermoneutral temperature (30°C) and treated with vehicle.

Cold exposure, compared to thermoneutral temperature, increased the total concentration of TAGs, as measured using conventional enzymatic assays (1.28±0.17 *vs*. 0.61±0.10 mmol/L, P=0.02), which was attenuated with Atglistatin treatment (0.75±0.22 mmol/L; P=0.90; **Supplementary figure 3A**). Moreover, cold exposure decreased 20/26 FFA species (77%, P<0.05) and increased 206/429 TAG species (48%, P<0.05; **Supplementary figure 3B**). The increase in TAGs after cold exposure included a range of TAG species with 0-8 double bonds and a total length from 46-56 carbons (**Figure 3A, B**). Atglistatin treatment completely blunted the cold-induced decrease in FFA lipid species, and largely attenuated the increase in TAG lipid species (**Supplementary figure 3C**). Specifically, with Atglistatin treatment cold exposure only increased 40/423 TAG species (9%, P<0.05) and decreased 13/423 TAG species (3%, P<0.05) (**Figure 3C, D**). The increase in TAGs and the decrease in FFAs were attenuated when mice were treated with Atglistatin and exposed to cold (**Supplementary Figure 3D**).

**Figure 3.**
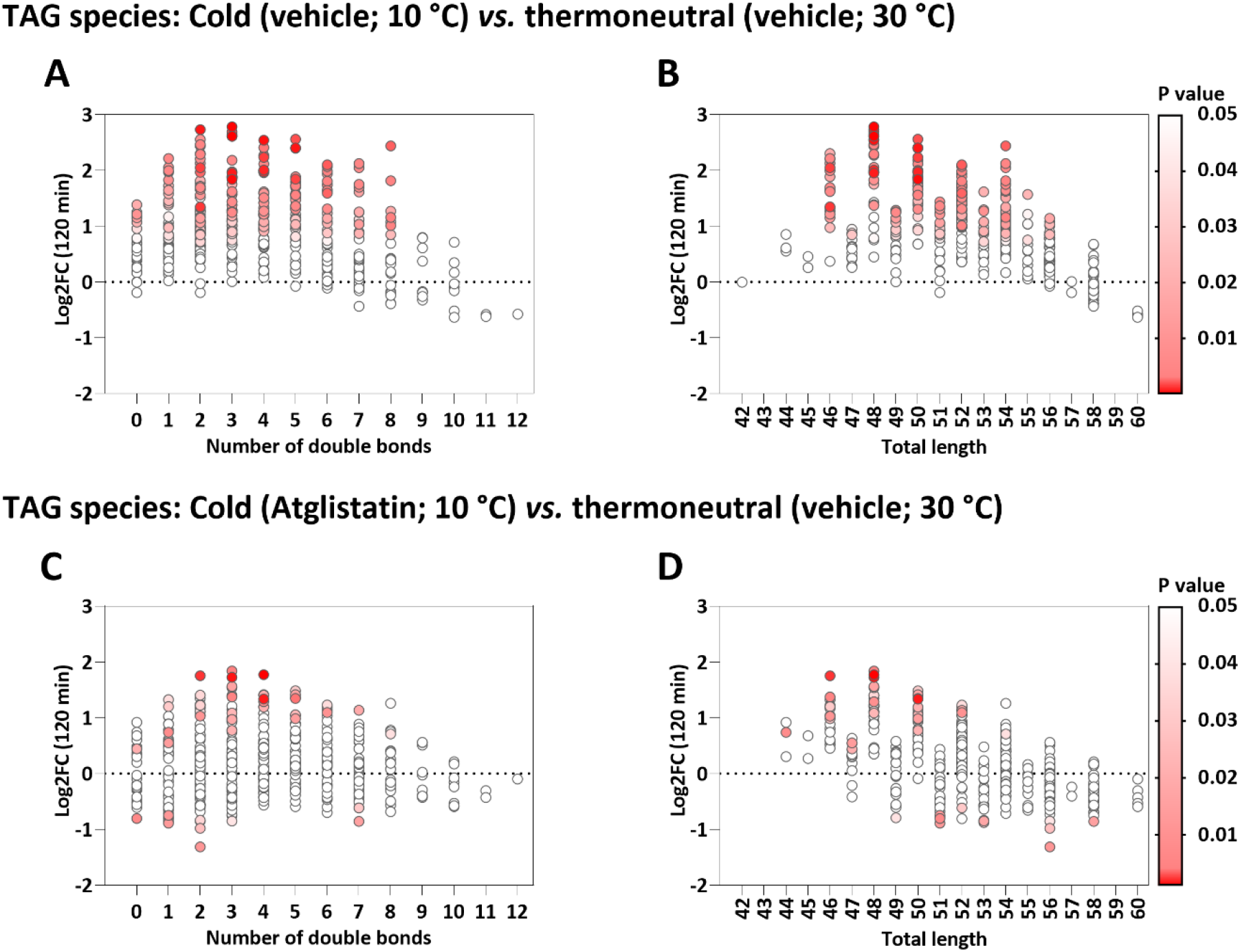
Triacylglycerol (TAG) species’ response to mild cold exposure in mice either treated with vehicle or with Atglistatin. (**A+B**) Bubble plots showing the P value (in color) and log2 fold change (log2FC; on y-axis) between mice exposed to cold and mice kept at thermoneutrality (n=6 each group; both treated with vehicle) per TAG species categorized by the number of double bonds (A) and total length (B). P values are obtained from paired Student’s t test. (**C+D**) Bubble plots showing the P value (in color) and log2FC (on y-axis) between mice exposed to cold and treated with Atglistatin and mice kept at thermoneutrality (n=6 each group; treated with vehicle) per TAG species categorized by the number of double bonds (C) and total length (D). P values are obtained from paired Student’s t test. FC, fold change; TAG, triacylglycerol.

## Discussion

In this study, we aimed to gain insight in the effect of cold exposure on TAG metabolism in humans by performing enzymatic assays combined with in-depth lipidomic profiling during cold exposure. We reveal that cold exposure increases a broad range of circulating FFAs and transiently decreases circulating total TAGs, accompanied with a dynamic change in circulating TAGs as dependent on their saturation status and length. During cold exposure, more saturated and shorter TAGs consistently decrease within 30 min, whereas TAGs enriched in PUFAs start to increase after 90-120 min. A mechanistic study in mice showed that the cold-induced increase in circulating TAGs is attenuated when blocking intracellular lipolysis. Our findings point towards a mechanism by which cold exposure causes avid intracellular lipolysis to liberate FFAs which stimulate the hepatic biosynthesis and release of PUFA-containing TAGs.

Until now, the effects of cold exposure on whole-body lipid metabolism in humans remained controversial, as some studies reported that cold exposure (for 180 minutes at 18°C) does not change total circulating TAG concentration^11,14^, whereas others even observed an increase in the total circulating TAG concentration when cold exposed for 120-150 minutes at shivering threshold^15,16^. In the current study, we observed a transient decrease in the total circulating TAG concentration at 30 min, an effect that was probably missed in those studies^11,14-16^. This was accompanied by a prolonged decrease in shorter, more saturated circulating TAG species. Mechanistically, cold exposure stimulates the sympathetic nervous system, resulting in the release of norepinephrine that activates adrenergic receptors on thermogenic tissues (*e*.*g*., skeletal muscle, brown adipose tissue)^22,23^. As a result, these tissues take up FAs from TAG-rich lipoproteins (*e*.*g*., very-low density lipoproteins, VLDL) as fuel for oxidation to eventually generate heat, hereby decreasing circulating TAGs^24,25^. Therefore, the VLDL-TAG-lowering effect of cold exposure as observed in mice^6-10^ can be recapitulated in humans.

After reaching a nadir at 30 min, prolonged cold exposure resulted in a gradual increase in circulating total TAGs, which was accompanied by an increase in longer, more unsaturated TAG species after 90-120 min of cold exposure. Mechanistically, such a gradual increase in total TAG concentration can be explained by an increased sympathetic outflow to the liver resulting in increased hepatic lipogenesis with subsequent increased release of TAG-containing VLDL particles into the circulation^15,26^. Interestingly, exposure to room temperature also continuously increased several TAG species, consistent with fasting-induced production of TAG-rich VLDL particles^27^. Considering that an initial decrease in circulating total TAGs and TAG species, as observed during cold exposure, was not observed during exposure to room temperature, probably indicates that the magnitude by which cold exposure induces uptake of TAG species by thermogenic tissues is underestimated.

Remarkably, during exposure to cold as well as room temperature, particularly PUFA-containing TAG species increased, although cold exposure increased most PUFA-TAG species (**Figure 2A** and **Supplementary figure 2**), suggesting that cold exposure enhances circulating PUFA-TAGs. The human body synthesizes most PUFAs from linoleic acid (LA, 18:2n6) and α-linolenic acid (ALA, 18:3n3), both of which are ‘essential’ PUFAs that need to be obtained from the diet. Biosynthesis of PUFAs is thereafter regulated through a complex step-wise series of reactions including both desaturation and elongation^28-30^. Desaturation in humans is regulated by delta-5 desaturase (encoded by *FADS1*) and delta-6 desaturase (encoded by *FADS2*)^28^, whereas elongation is regulated by elongases ‘Elongation of very-long-chain fatty acids 2 and 5’ (encoded by *ELOVL2* and *ELOVL5*) expressed in liver and other metabolic tissues^31-33^. Only a single previous study has reported on cold-induced lipidomic changes in humans. Using a cooling vest at 14°C for 1 h, only a few circulating FFA and TAG species were modulated, independent of the length or saturation status of the lipid species^34^. The seeming discrepancy with our results can easily be explained by the milder and shorter nature of the cooling protocol they used compared to ours, as we observe the increase in PUFA-containing TAGs at the end of cold exposure. Taken together, it is thus reasonable to assume that prolonged cold exposure increases elongation and desaturation.

In mice, the remodeling of specific TAG lipid species in metabolic tissues after long-term cold exposure has been demonstrated before^35-37^. That is, 24 hours of cold exposure at 5°C was shown to relatively increase PUFA-containing TAGs in the liver while uniformly decreasing TAG lipid species in brown adipose tissue^36^, suggesting that increased hepatic biosynthesis of TAGs is important to sufficiently refuel metabolic tissues with FAs during a strong cold stimulus. On the other hand, 3 days of cold exposure (4°C) is reported to cause an accumulation in long chain- and very-long chain-TAGs in murine white^37^ and brown adipose tissue^35^, most likely due to activated FA elongation pathways after long-term cold exposure. Collectively, we propose that the increase in circulating PUFA-containing TAGs after short-term cold exposure observed in our study is the result of their biosynthesis by the liver^28-30^, that is enhanced by cold exposure.

Cold exposure also causes increased sympathetic activation of white adipose tissue to increase intracellular lipolysis, resulting in an increased flux of FFAs toward the liver^38^, which enhances the biosynthesis and release of TAG-rich VLDL particles into the circulation^39,40^. To evaluate whether intracellular lipolysis is crucial for the cold-induced increase in TAG species, we performed a mechanistic study assessing the effect of blocking ATGL by Atglistatin during cold exposure. ATGL is particularly highly expressed in adipose tissue, and to a lower extent in other tissues as the liver and skeletal muscle^41,42^, and is important for mobilizing lipids by catalyzing the first step in TAG lipolysis^42^. Indeed, the increase in total TAG concentration and many TAG species as induced by cold exposure was abolished by Atglistatin, suggesting crucial involvement of FFA-driven hepatic TAG production. Provided these data can be translated to humans, the cold-induced increase in circulating TAG species is dependent on intracellular lipolysis in white adipose tissue.

The cold-activated dynamic changes in lipids as shown here help uncover mechanisms by which the application of cold exposure can benefit cardiometabolic health. In obesity, white adipose tissue is key in the development of metabolic disorders by spillover of FFA induced by insulin resistance, leading to elevated circulating FFAs^43^ that cause ectopic lipid deposition in organs including the liver^44,45^. Our present data suggest that cold exposure may counteract ectopic lipid deposition by increased uptake by thermogenic tissues as fuel for oxidation. Interestingly, in mice dietary (omega-3) PUFAs trigger thermogenesis by brown adipose tissue^46^, probably by sympathetic activation^47^, or through the G-protein-coupled receptor GPR120^48^. In addition, it has become increasingly clear that enhancing circulating PUFAs by dietary intake of (omega-3) PUFAs is a promising way to improve cardiometabolic health, due to their anti-inflammatory and TAG-lowering effects^49-52^. Similarly, PUFAs are precursors of oxylipins and fatty acid esters of hydroxy fatty acids (FAHFAs), which significantly increase in the circulation after cold exposure. The cold-induced biosynthesis of PUFA-TAG species may thus promote thermogenic processes and/or mediate anti-inflammatory responses, warranting further studies to fully elucidate the role of PUFAs in the thermogenic function, as well as their relevance for improving metabolic health in obesity.

This work is not without limitations. Although it is quite unlikely that it has impacted the results, the control condition was systematically performed 2 hours later than the experimental condition. The lipidomic profile has been assessed in blood samples only, therefore, the proposed mechanisms remain speculative and should be investigated further, preferably by repeated invasive tissue biopsies taken during cold exposure. In addition, we identified lipids on a species level (*e*.*g*., TAG 56:8-FA 16:1), which provides information on one out of three individual FA chains. Consequently, for two out of three FA chains omega-3 and omega-6 PUFAs cannot be distinguished. Furthermore, we used the ATGL inhibitor Atglistatin to elucidate the importance of intracellular lipolysis on the cold-induced increase in circulating TAGs. It is important to note that ATGL is also present in other tissues than white adipose tissue^41,42,53^, as a result of which it is possible that FAs for TAG synthesis are derived from non-adipose tissues. Future studies addressing cold-induced lipidomic profiles in larger and more diverse populations (*e*.*g*., women, people with insulin resistance or obesity) are required to further address utility of cold exposure in metabolic health.

In conclusion, short term cold exposure of young men increases circulating FFAs, accompanied by a decrease in circulating shorter, more saturated TAGs, and a subsequent increase in longer, PUFA-enriched TAGs. Combined with a mechanistic study in mice, these findings are consistent with a mechanism in which cold exposure increases lipolysis in white fat to drive hepatic TAG production to feed thermogenic tissues (for graphic summary, see **Figure 4**). The discovery that cold exposure elicits modifications in lipid composition including an increase in PUFA-containing TAGs sheds new light on the beneficial effects of cold exposure and warrants future studies to determine the potential contribution of *e*.*g*. long-term repetitive cooling to cardiometabolic health.

**Figure 4.**
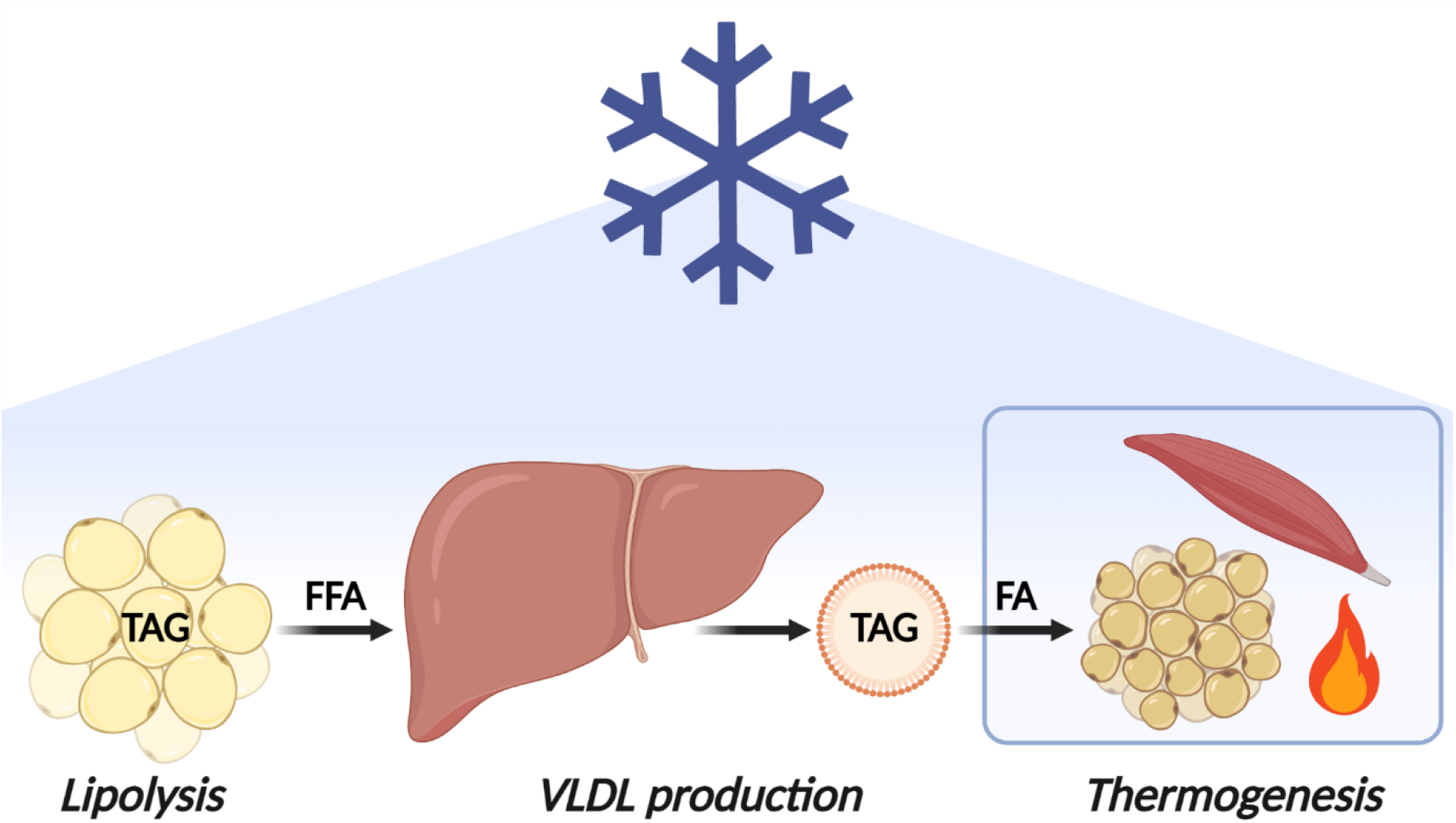
Graphic summary of proposed mechanism. We propose a mechanism by which cold exposure induces intracellular lipolysis in white adipose tissue, increasing circulating free fatty acids that can subsequently drive hepatic biosynthesis and release of triacylglycerol (TAG)-containing very-low density lipoprotein (VLDL) particles to feed thermogenic tissues. The cold-activated thermogenic tissues (*i*.*e*., brown adipose tissue and skeletal muscle) take up TAG-derived fatty acids from VLDL particles as fuel for oxidation to generate heat. FA, fatty acid; FFA, free fatty acids; PUFA, polyunsaturated fatty acid; TAG, triacylglycerol; VLDL, very-low density lipoprotein.

## Database and data handling procedures

**The study has a data management plan that strictly follows the regulation of the Leiden University Medical Center (LUMC), which ensures data protection all the project. The data has been treated according to EU regulation 2016/679 (e.g**., **pseudo anonymization, security measures to prevent unauthorized access). All the data and research material were stored following the guidelines of LUMC Biobank Endocriene Ziekten. Our data management plan also followed the H2020 FAIR principle, as we were making our research data findable, accessible, interoperable (i.e**., **allowing data exchange between researchers) and reusable (while having in mind the embargo policies)**.

## Author contributions

Conceptualization, S.K., P.C.N.R. and B.M.T.; Methodology, M.G, P.C.N.R. and B.M.T.; Validation, M.G., G.F.G. and R.Z.; Formal Analysis, M.E.S., L.J.F. and B.M.T.; Data collection, K.J.N., L.J. and Z.Y.; Data Curation, M.E.S., L.J.F. and B.M.T.; Writing – Original Draft, M.E.S. and L.J.S.; Writing – Review & Editing, P.C.N.R. and B.M.T.; Visualization, M.E.S. and L.J.S.; Supervision, M.R.B., P.C.N.R. and B.M.T.; M.E.S. and L.J.F. share first authorship. The order in which they are listed reflects the workload from initiation of the study until publication. All authors commented on the manuscript and approved the final version of the manuscript.

## Data sharing

The data that support the findings of this study are available from the corresponding author upon reasonable request, as the study consists of a high number of participants and outcomes and requires specific knowledge for data interpretation.

## Acknowledgements

We thank Niek Blomberg for his excellent support with the lipidomics analyses, and Trea Streefland for her technical assistance with the measurement of FFAs and TAGs using enzymatic kits. The graphical summary figure (**Figure 4**) was created with BioRender.com.

