## Supplementary table for "Cold exposure induces dynamic changes in circulating triacylglycerol species, which is dependent on intracellular lipolysis: a randomized cross-over trial"

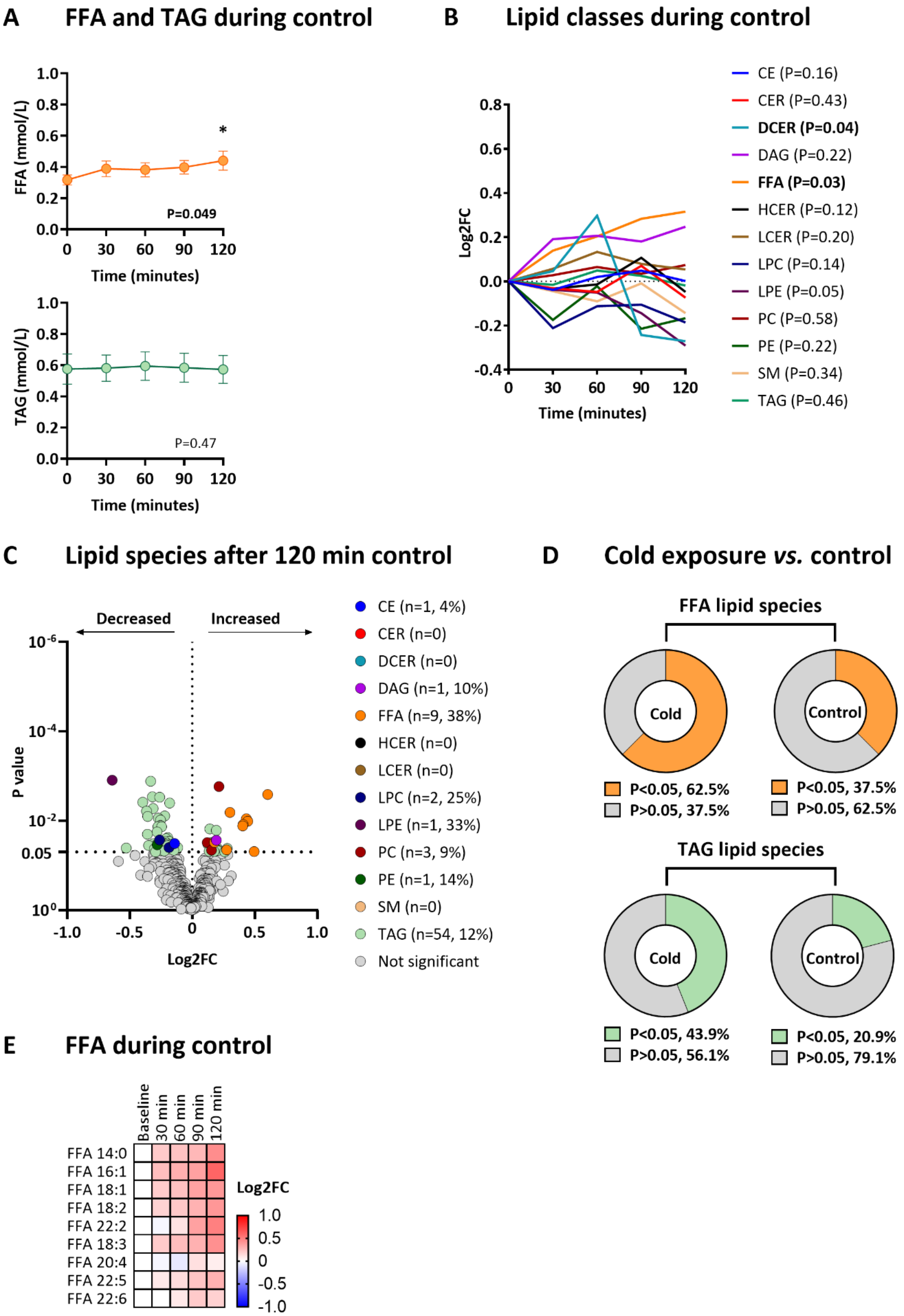
A
FFA and TAG during control
FFA (mmol/L)
P=0.049
*
TAG (mmol/L)
P=0.47
Time (minutes)
B
Lipid classes during control
Log2FC
Time (minutes)
CE (P=0.16)
CER (P=0.43)
DCER (P=0.04)
DAG (P=0.22)
FFA (P=0.03)
HCER (P=0.12)
LCER (P=0.20)
LPC (P=0.14)
LPE (P=0.05)
PC (P=0.58)
PE (P=0.22)
SM (P=0.34)
TAG (P=0.46)
C
Lipid species after 120 min control
Decreased
Increased
P value
Log2FC
CE (n=1, 4%)
CER (n=0)
DCER (n=0)
DAG (n=1, 10%)
FFA (n=9, 38%)
HCER (n=0)
LCER (n=0)
LPC (n=2, 25%)
LPE (n=1, 33%)
PC (n=3, 9%)
PE (n=1, 14%)
SM (n=0)
TAG (n=54, 12%)
Not significant
D
Cold exposure vs. control
FFA lipid species
Cold
Control
P<0.05, 62.5%
P>0.05, 37.5%
P<0.05, 37.5%
P>0.05, 62.5%
TAG lipid species
Cold
Control
P<0.05, 43.9%
P>0.05, 56.1%
P<0.05, 20.9%
P>0.05, 79.1%
E
FFA during control
Baseline
30 min
60 min
90 min
120 min
FFA 14:0
FFA 16:1
FFA 18:1
FFA 18:2
FFA 22:2
FFA 18:3
FFA 20:4
FFA 22:5
FFA 22:6
Log2FC

**Figure S1. Lipidomic changes during the control experiment without personalized cold exposure**

(**A**) Dynamic changes in total free fatty acid and triacylglycerol concentration during the control experiment, as measured with enzymatic assays. Data is presented as mean and standard error of the mean. P values are obtained from ANOVA repeated measures. *P<0.05; compared to baseline (*i.e.*, 0 min). (**B**) Dynamic changes in lipid classes during the control experiment. Data is presented as the log2 fold change (log2FC) relative to baseline (*i.e.*, 0 min). ANOVA repeated measures were used to compare the different timepoints. (**C**) Volcano plot showing the change of individual lipid species after 120 min during the control experiment exposed to room temperature. The X-axis shows the log2FC between 120 min *vs.* baseline, the Y-axis shows the P value. P values are obtained from paired Student's t test. Values between brackets indicate the absolute number and the percentage of lipid species within the lipid class that were modulated. (**D**) FFA and TAG species that significantly dynamically changed during cold exposure (left) or during the control experiment (right). P values are obtained from ANOVA repeated measures, percentages indicate the lipid species within the lipid class that were modulated. (**E**) Heatmap of FFA species that significantly dynamically changed during the control experiment. The color of the squares represents the log2FC relative to baseline. FFAs are sorted by the number of double bonds. ANOVA repeated measures were used to compare the different timepoints. CE, cholesteryl esters; CER, ceramides; DAG, diacylglycerols; DCER, dihydroceramides; FFA, free fatty acids; FC, fold change; HCER, hexosylceramides; LCER, lactosylceramides; LPC, lysophosphatidylcholine; LPE, lysophosphatidylethanolamine; PC, phosphatidylcholine; PE, phosphatidylethanolamine; SM, sphingomyelin; TAG, triacylglycerols.

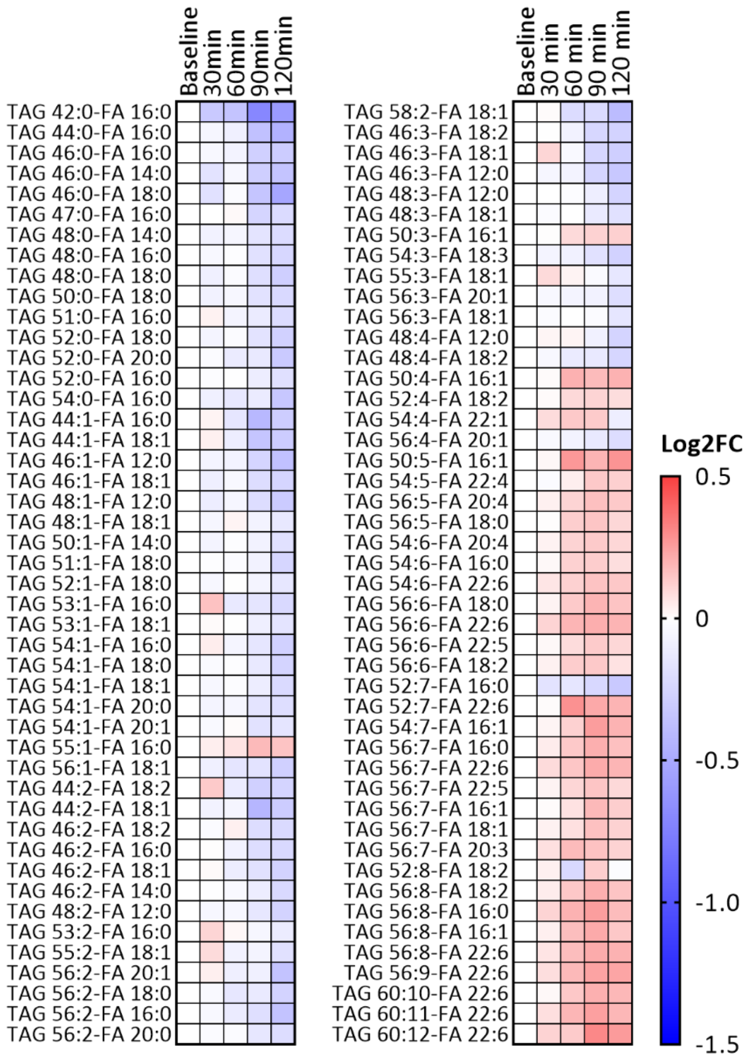


**Figure S2. Triacylglycerol (TAG) species during the control experiment without personalized cold exposure**

Heatmap of TAG species that significantly dynamically changed during the control experiment. The color of the squares represents the log2 fold change (log2FC) relative to baseline. TAGs are sorted by the number of double bonds. ANOVA repeated measures were used to compare the different timepoints. FA, Fatty acid; FC, fold change; TAG, triacylglycerol.

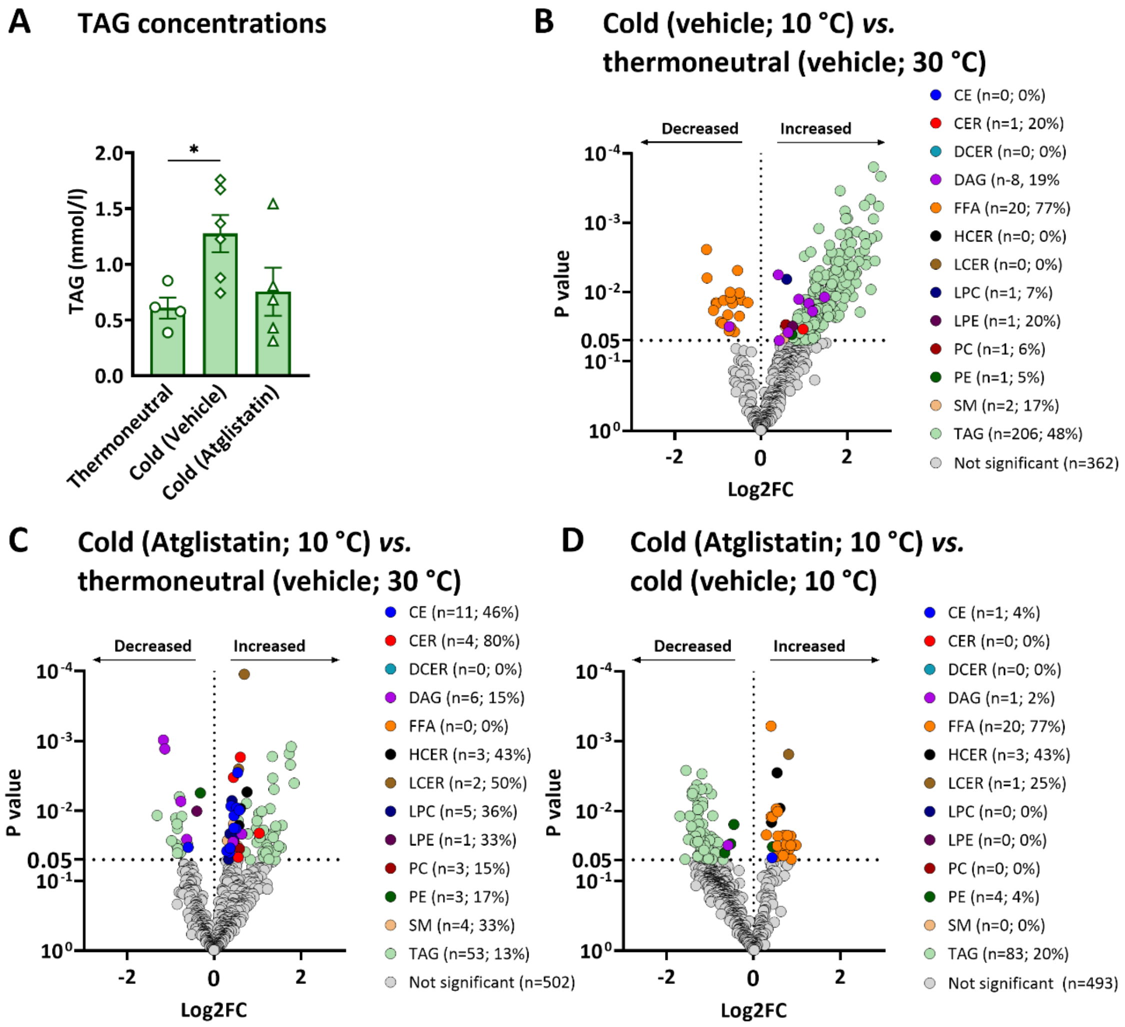


**Figure S3. The effect of cold exposure on individual lipid species in mice**

(**A**) Total triacylglycerol (TAG) concentration of mice just after the intervention, as measured with an enzymatic assay. Data is presented as mean and standard error of the mean. P value is obtained from Mann Whitney U test. (**B-D**) Volcano plot showing the differences in lipid species between mice exposed to cold *vs.* mice kept at thermoneutrality (B) (n=6 each group; both treated with vehicle); between mice exposed to cold (n=6; treated with Atglistatin) *vs.* mice kept at thermoneutrality (treated with vehicle) (C); and between mice exposed to cold (n=6; treated with Atglistatin) *vs.* mice exposed to cold (n=6; treated with vehicle) (D).The X-axis shows the log2 fold change (log2FC) between the groups, the Y-axis shows the P value. P values are obtained from unpaired Student's t test. Values between brackets indicate the absolute number and the percentage of lipid species within the lipid class that were modulated by cold exposure (B, C), and by Atglistatin treatment (D). CE, cholesteryl esters; CER, ceramides; DAG, diacylglycerols; DCER, dihydroceramides; FFA, free fatty acids; FC, fold change; HCER, hexosylceramides; LCER, lactosylceramides; LPC, lysophosphatidylcholine; LPE, lysophosphatidylethanolamine; PC, phosphatidylcholine; PE, phosphatidylethanolamine; SM, sphingomyelin; TAG, triacylglycerols.
